# Discriminative and affective touch converge: Somatosensory cortex represents Aß input in a CT-like manner

**DOI:** 10.1101/2021.07.21.453292

**Authors:** Annett Schirmer, Oscar Lai, Francis McGlone, Clare Cham, Darwin Lau

## Abstract

Current theory divides the human mechanical sense into discriminative and affective systems. A discriminative system supports tactile exploration and manipulation via fast Aß signaling, whereas an affective system supports the pleasure of friendly interpersonal touch via slow CT signaling. To probe this system segregation, we recorded the electroencephalogram from participants being stroked and reporting stroke pleasantness. We observed a somatosensory negativity that was maximal for CT optimal as compared with sub-optimal velocities, that predicted subjective pleasantness, and showed only for stroking of hairy skin known to be CT innervated. Importantly, the latency of this negativity preceded C fiber input to the brain by several hundred milliseconds and is best explained by interactions between CT and Aß processes in the spinal cord. Our data challenge the divide between discriminative and affective touch implying instead that both fast Aß and slow CT signaling play an important role in tactile pleasure.

## Introduction

To humans, as to most other mammals, physical contact with conspecifics can be rewarding. In this context, low pressure gentle stroking appears to be a particularly potent stimulus that excites privileged processing pathways supported by a special C-fiber mechanoreceptor referred to as C-tactile (CT) afferent (Löken et al., 2009; Olausson et al., 2002; Vallbo et al., 1999; Vallbo and Johansson, 1984). Prior research has helped characterize the peripheral physiology of this receptor and shed light on potential projection targets in the brain. However, the mechanisms by which CT activation induces pleasure are still unknown and the extent to which other receptor types may be relevant remains debated. Here, we sought to tackle these issues by using the electroencephalogram (EEG) to examine possible cortical CT correlates and how they unfold in time.

Current thinking, based primarily on peripheral nerve anatomy, divides the human sense of touch into an affective and a discriminative system (Case et al., 2016; McGlone et al., 2014). Affective touch serves homeostatic and social functions. It has been linked to the above-mentioned C fibers, which are unmyelinated and therefore have a slow conduction velocity. While non-tactile C fibers such as those supporting the perception of pain, itch, or temperature are known to project via the spinothalamic tract, CT spinal projection pathways are still incompletely documented and their inclusion in the spinothalamic tract remains debated (Marshall et al., 2019). It is known, however, that their input reaches the insula directly via the thalamus and by-passing somatosensory cortex (Olausson et al., 2002).

Discriminative touch helps us detect, within tens of milliseconds, any physical stimulus on the body surface and, via haptic mechanisms, to identify and manipulate objects and tools. Discriminative touch receptors include a variety of Aß low threshold mechanoreceptors (LTMRs) innervated by myelinated, and therefore fast conducting, nerves. First-order LTMR neurons synapse in the dorsal horn of the spinal cord, from where second order neurons travel via the dorsal-column medial lemniscus pathway to the thalamus and from there to primary and secondary somatosensory cortex in the postcentral gyrus (McGlone et al., 2014).

A primary reason for the functional dissociation between CT and Aß signaling is that they differ in how they respond to changes in the velocity of a stroking stimulus. CTs show an inverted u-shaped relationship between firing rate and stroking velocity that peaks for velocities from 1 to 10 cm/s (Löken et al., 2009). Similarly, subjective feelings of tactile pleasure peak for velocities from 1 to 10 cm/s and are lower for faster and slower stroking (Essick et al., 1999). Thus, CT firing linearly predicts tactile pleasure (Löken et al., 2009). In contrast, Aß receptors fire in a monotonically increasing manner with increasing stroking velocity (Löken et al., 2009). Their firing seems unrelated to tactile pleasure, but may instead be relevant for sensory properties such as touch intensity (Case et al., 2016). Additionally, initial evidence implies that Aß activity is less pleasurable than CT activity (Löken et al., 2009).

Despite the theoretical divide between affective and discriminative touch, there is some empirical data suggesting that perhaps both systems support tactile pleasure and could serve homeostatic and social functions. For one, recent psychophysical research found that touch to glabrous and hairy skin can be equally pleasurable (Luong et al., 2017; Pawling et al., 2017; Schirmer and Gunter, 2017). CT afferents, as defined by their velocity response curve, have been documented only in hairy skin. Moreover, a recent report of LTMRs in palmar skin demonstrated only sparse innervation (one unit each was found in 3 out of 43 participants, only one of these units was located in skin used for grasping) (Watkins et al., 2020). By contrast, Aß fibers are found across skin types and are particularly dense in the palm. Thus, Aß and CT fibers may be similarly relevant for tactile pleasure.

Additionally, there is evidence for interactions between Aß and C fiber processes. Such evidence was first reported in the context of C fiber pain signaling and inspired what is known as the gate control theory of pain (Melzack and Wall, 1965). According to this theory, Aß signals arriving in the dorsal horn can dampen the onward projection of C signals through the activity of interneurons. Despite some conflicting data (e.g., Nathan and Rudge, 1974), the theory has been largely supported and remains relevant in the field (Mendell, 2014).

In line with the gate control theory, there is now evidence from non-human animals indicating that non-painful tactile transmission in the dorsal horn LTMR recipient zone integrates input from different fiber types. Recent cellular work in rodents revealed a convergence of Aß and CT input onto overlapping interneurons indicating that faster Aß input could modulate the onward projection of slower CT input and vice versa (Abraira et al., 2017).

Importantly, however, whether Aß and CT pathways interact in humans and how their interaction might impact cerebral tactile processing is still unknown. To date, the human literature has focused on measuring peripheral nerve activity (Löken et al., 2009; Watkins et al., 2020, 2017) and pursued brain processes with techniques affording good spatial, but poor temporal resolution (Case et al., 2016; Morrison, 2016; Olausson et al., 2002). Indeed, we know of only three studies examining the time course of forebrain cortical processes.

One EEG study delivered strokes at CT optimal velocities and tracked a stimulus associated response referred to as an ultra-late positive potential (Ackerley et al., 2013). The authors suggested that CTs modulated this potential at a frontal midline region (Fz) about 700 ms following stimulus onset. As CT input tends to travel at ∼1 m/s (Vallbo et al., 1993; Watkins et al., 2017, p. 201), the arrival of signals from the arm at the brain was estimated to take about 700 ms. A subsequent MEG study also identified a signal modulation around 700 ms following CT targeted touch and identified sources in the posterior insula (Hagberg et al., 2019). Last, an EEG study presented a 300 ms stimulus at 30 cm/s and a 3 s stimulus at 3 cm/s targeted at Aß and CTs, respectively (Haggarty et al., 2020). The results failed to replicate the frontal midline effect at 700 ms and instead showed a positive potential for both conditions at the time stimulus offset. No study reported CT effects over somatosensory cortex.

Here, we set out to extend earlier work and to scrutinize CT and Aß signaling in the brain by leveraging on the high temporal resolution of the EEG. Specifically, we conducted two experiments manipulating stroking velocity and skin type to establish fiber-specific as well as fiber-integrative processes. In the first experiment, we delivered a stroking stimulus at CT optimal and sub-optimal speeds to the hairy forearm, which is innervated by both CT and Aß fibers. After each stroke, participants rated the pleasantness of the sensation. We examined Aß signaling between 0 to 700 ms following touch onset over somatosensory leads. Additionally, we pursued CT signaling after 700 ms over a frontal midline electrode. The second experiment shifted touch from the arm to the palm as to ascertain a role of CT afferents in the observed effects.

Our experiments probed the following two hypotheses. First, if frontal midline cortex presents CT relevant processes as suggested previously (Ackerley et al., 2013), we expected to see a positive ERP 700 ms after stimulus onset with an amplitude that tracks velocity in an inverted u-shaped manner for input from hairy but not glabrous skin. Second, and more importantly, we speculated that somatosensory cortex receives CT-relevant input earlier, via Aß pathways that are modulated by CT signals in the spinal cord. If so, ERPs recorded over somatosensory cortex might show a similar velocity dependence as frontal midline ERPs. Moreover, they should do so within 700 ms of touch onset for hairy, but not glabrous skin.

### Results of Experiment 1 – Stroking of Hairy Skin

#### ERPs

In line with a previous report (Ackerley et al., 2013), visual inspection of the 4 s ERP epochs at Fz revealed a positive deflection starting around 700 ms after touch onset. This modulation lasted for about 300 ms after which there were no further visible effects (Figure 1A). Because long ERP epochs are associated with lower trial numbers following artifact rejection, we re-analyzed our EEG data with shorter epochs as described in the methods. We then conducted our statistical analysis on Fz voltages in keeping with an earlier study (Ackerley et al., 2013).

**Figure 1.**
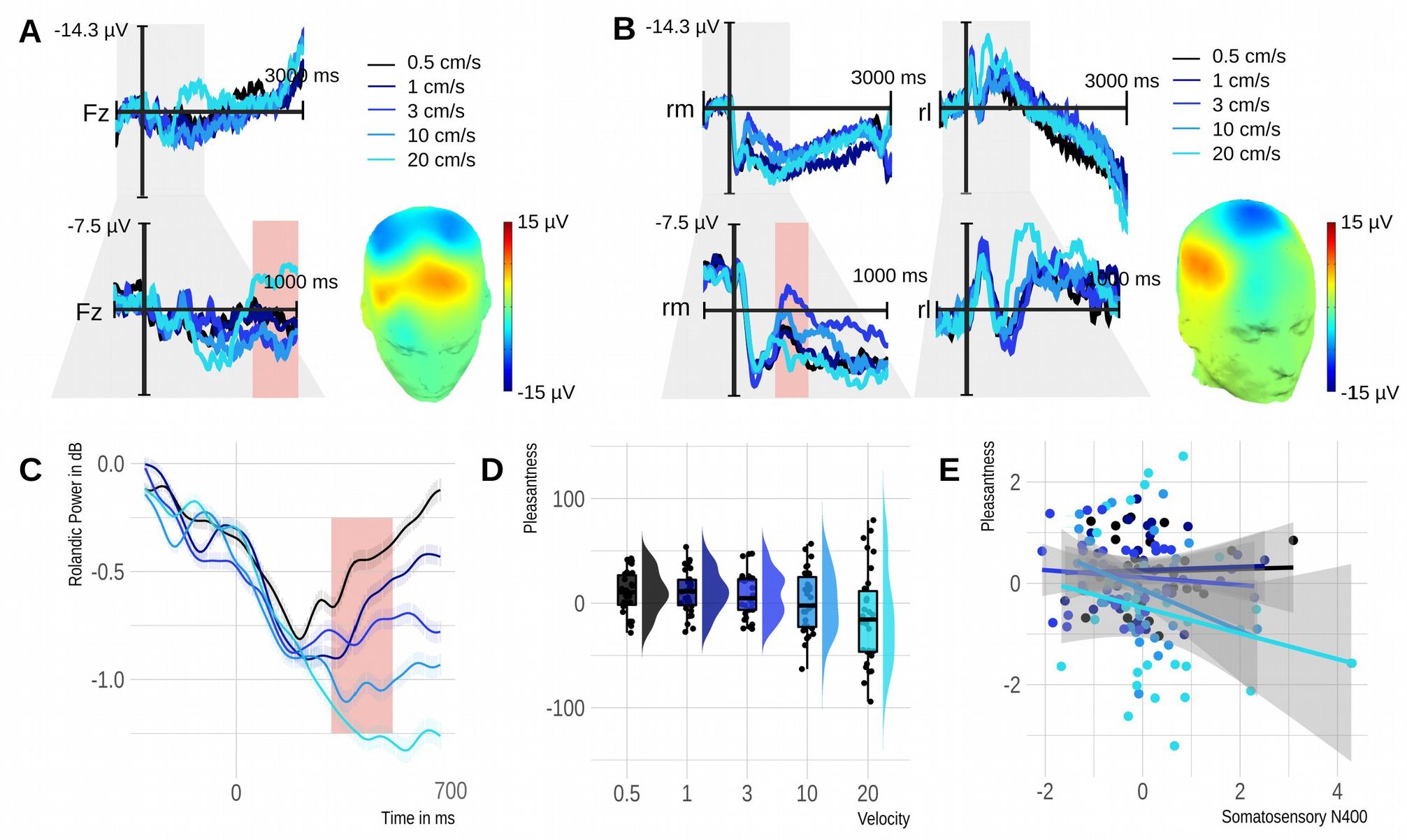
Results of Experiment 1. A) Velocity effect observed at Fz, a site previously linked to an ultra-late positive potential. B) Velocity effect observed over the somatosensory region contra-lateral to the tactile stimulus which we refer to as somatosensory N400. Reddish rectangle highlights windows for statistical analysis ranging from 0.7 to 1 seconds for Fz and from 0.3 to 0.5 seconds for the somatosensory ERP and Rolandic power. The topography of each velocity effect (3 cm/s minus 20 cm/s) is printed on a 3D head. Fz – frontal midline electrode, rm – right medial region of interest (Cz, C2, CP2), rl – right lateral region of interest (C6, T8, CP6). C) Time course of power changes evoked by tactile stimuli in alpha and beta bands. D) Rain plots illustrate a change in subjective pleasantness as a function of stimulus velocity. E) Scatter plots with regression lines showing the linear relationship between the normalized subjective pleasantness and the normalized N400 amplitude at the subject level. Overall, larger N400 amplitudes (i.e., more negative voltages) were associated with greater pleasure.

Fz mean voltages from 700 to 1000 ms entered a mixed effects model with the second order polynomial of the natural velocity logarithm as the fixed effect and the participants’ intercept as the random effect. For this and the other brain measures, we could not model the participants’ slope as we had only one value per participant and condition. F-statistics were obtained using the Satterthwaite approximation for degrees of freedom. Our analysis returned a significant velocity effect (F[2,118]=5.23, p=.007). Additionally, a likelihood ratio test including a model without the polynomial indicated that the polynomial offered the better fit (X[1]=18.06, p=.007).

Apart from the Fz effect, we observed an interesting negative deflection over somatosensory cortex contra-laterally to the stimulation, which we refer to as a somatosensory N400. Like the Fz effect, the N400 showed a velocity-dependent negative quadratic pattern but emerged significantly earlier between 300 to 500 ms following touch onset. For analysis, we computed mean voltages within this time window and across a subset of right centro-medial electrodes (Cz, C2, CP2). Please refer to Figure 1B which illustrates the dipole of this effect and guided electrode selection (the negativity peaked centrally and reversed polarity laterally). The polynomial mixed effects model was again significant (F[2,118]=13.17, p<.001) and was superior when compared with the simple linear model using a likelihood ratio test (X[1]=32.17, p<.0001).

#### Rolandic rhythms over contra-lateral somatosensory cortex

Given the novelty of the observed N400 effect, we examined whether overall activity in somatosensory cortex increased with increasing velocity as one would expect for Aß input (Case et al., 2016; Löken et al., 2009). Like the ERP, mean power between 300 and 500 ms following touch onset in the alpha and beta band at right centro-medial electrodes was subjected to a polynomial mixed effects model, which revealed statistically significant results (F[2,118]=24.73, p<.001). Note, however, that, as predicted, the quadratic term indexed a monotonical decrease in power with increasing velocity (Figure 1C). Again, the polynomial fit was superior to a linear fit (X[1]=5.32, p=.021).

#### Pleasantness Ratings

Rating data were analyzed using a similar statistical approach as for the EEG data. Trial-wise rating scores were subjected to a model with the second order polynomial of the natural velocity logarithm as the fixed effect and the participants’ intercepts and slopes as the random effects. This revealed a significant velocity effect (F[2,29]=3.47, p=.044). Additionally, a likelihood ratio test comparing this original model against one with velocity modeled linearly revealed that the former was a significantly better fit (X[4]=879.61, p<.0001).

Last, we examined which of the above-mentioned brain measures predict pleasantness ratings. Mean pleasantness served as the dependent variable, a given brain measure and velocity served as fixed effects, and the participants’ intercepts served as the random effect. Velocity was included simply to reduce Type 1 error and was of no interest. For the Fz ERP, the result was non-significant (p=.367). However, we observed a significant relationship for the somatosensory N400 (F[1,117]=4.21, p=.042) indicating that more negative ERPs were associated with greater pleasantness. The power of Rolandic rhythms over the somatosensory region was unrelated with pleasantness (p=.742).

### Results of Experiment 2 – Stroking of Glabrous Skin

#### ERP

Again, we first explored a late frontal midline modulation between 700 and 1000 ms (Figure 2A). As for hairy skin, our statistical analysis returned a significant quadratic effect of velocity on the mean ERP amplitude (F[2,118]=4.38, p=.015). Mid-range velocities elicited a more positive ERP deflection than the slowest and fastest velocities. A likelihood ratio test identified the quadratic model as superior to a linear one (X[1]=16.58, p<.0001).

**Figure 2.**
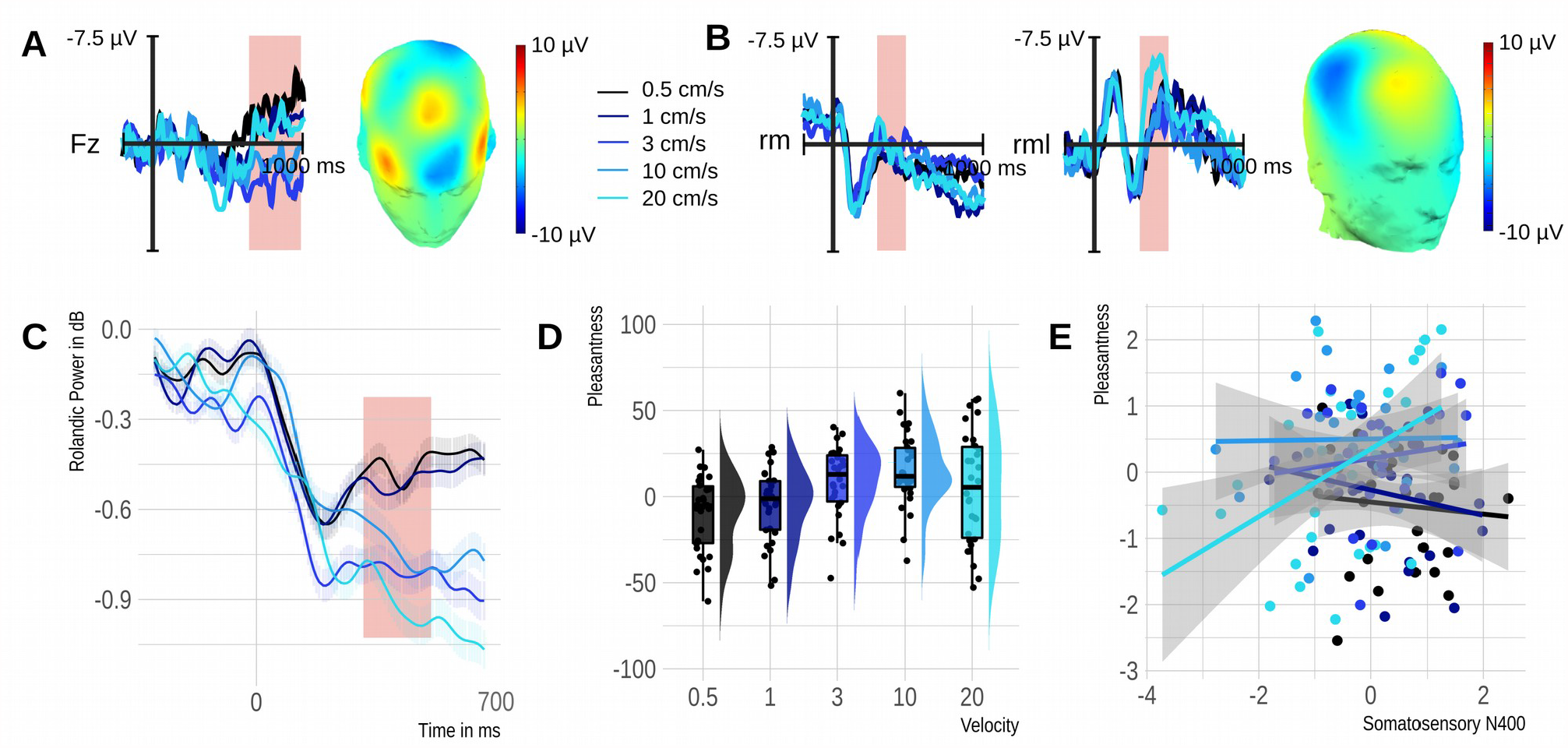
Results of Experiment 2. A) Velocity effect observed at Fz. B) Somatosensory N400 velocity effect over the right medial region explored in Experiment 1 (rm; Cz, C2, CP2) and slightly more lateral region adapted for the cortical representation of the palm (rml; C2, C4, CP4). Reddish rectangles highlight windows for statistical analysis ranging from 0.7 to 1 seconds for the Fz effect and from 0.3 to 0.5 seconds for the somatosensory N400 and Rolandic power. The topography of each velocity effect (Fz - 3 cm/s minus 20 cm/s; rml – 20 cm/s minus 0.5 cm/s) is printed on a 3D head. Fz – frontal midline electrode, rm – right medial region of interest, rml – right medial/lateral region shifted to accommodate the more lateral representation of the palm. C) Time course of power changes evoked by tactile stimuli in alpha and beta bands. D) Rain plots illustrate a change in subjective pleasantness as a function of stimulus velocity. E) Scatter plots with regression lines showing the absence of a linear relationship between the normalized subjective pleasantness and the normalized N400 amplitude at the subject level.

Next, we examined the somatosensory N400 as identified in Experiment 1. The result was non-significant (p=.614). Because the hand of the somatosensory homunculus sits a bit more laterally than the forearm, we also examined a more lateral region comprised of C2, C4, and CP4. Here, a statistical analysis was significant (F[2,118]=15.59, p<.001). Moreover, a quadratic model was again a better fit than a linear model (X[1]=8.51, p<.001). Importantly, however, rather than indexing an inverted u-shape, it indexed a monotonic increase in negativity with increasing velocity. The N400 was larger for the fastest compared with the slower velocities (Figure 2B).

#### Rolandic rhythms over contra-lateral somatosensory cortex

As for the ERP, we examined the power of Rolandic rhythms for the original somatosensory region identified in Experiment 1 and for a more lateral one. Over the original region, the velocity main effect was significant (F[2,118]=8.88, p<.001) and the quadratic fit was better than the linear fit (X[1]=6.24, p=.012). These results replicated over the more lateral region (F[2,118]=9.38, p<.001; X[1]=4.63, p=.031) and reflected a monotonical decrease in power with increasing velocity (Figure 2C).

#### Pleasantness Ratings

Analysis of rating data revealed a main effect of velocity indicating that its quadratic effect on Pleasantness was significant (F[2,29]=9.94, p<.001). Additionally, the likelihood ratio test demonstrated that the quadratic fit was better than a linear fit (X[4]=1462,p<.0001).

Last, we tested whether, as in Experiment 1, ERP and Rolandic rhythms could predict participants’ pleasantness ratings. Again, there was no effect for the frontal midline ERP (p=.729) and the somatosensory N400 over the original statistical region was no longer significant (p=.315). Moreover, there was only a marginal effect for the N400 over the more lateral somatosensory region (F[1,103]=2.89, p=.092) and the effect was opposite to that of Experiment 1 and its replication (Figure 2D). Power of Rolandic rhythms over the original and the more lateral somatosensory region was unrelated to pleasantness (ps>.46).

## Discussion

The present study probed whether a frontal midline effect reflects CT signaling and examined somatosensory brain potentials for possible evidence of Aß and CT interactions.

In line with earlier research (Ackerley et al., 2013), a frontal midline effect emerged consistently at ∼700 ms following touch onset. Moreover, as predicted, its associated positive deflection tracked stroking velocity. CT optimal stimulus velocities elicited more positive voltages than faster and slower velocities in line with the idea that CT input modulates frontal midline processes. Notably, however, other aspects of our results contradicted a CT effect. For one, the frontal midline ERP was comparable for hairy and glabrous skin and as such seemed independent of CT activity. Second, this positivity failed to statistically predict tactile pleasantness at the subject level.

In light of these results, we reconsider the functional role of frontal midline responses to CT-targeted touch. Rather than being driven strictly by CTs, they must reflect a more general, pathway independent cortical representation of tactile input. This conclusion agrees with past research demonstrating an overlap in higher-order perceptual processes for hairy and glabrous touch (Luong et al., 2017; Pawling et al., 2017). It also converges with findings that apart from pleasure, other psychological properties show a bias for CT optimal velocities including perceived human-likeness (Wijaya et al., 2020) and smoothness (Sailer et al., 2020). Thus, we reason that frontal midline responses are removed from basic sensory processes and support, perhaps, a combination of the subjective feeling aspects arising from touch.

Examination of somatosensory brain potentials revealed the expected evidence for overlap between putative discriminative and affective touch pathways. Specifically, it identified a novel ERP response that peaked at 400 ms following touch onset over somatosensory cortex contra-lateral to the stimulation site. This response was characterized by a negative deflection, which we refer to as a somatosensory N400, and tracked stroking velocity in a CT-like manner. Participants in two independent samples who were stroked on the forearm displayed larger N400 amplitudes for CT optimal velocities when compared with faster and slower velocities and, across conditions, N400 amplitudes statistically predicted the participants’ mean pleasantness ratings. Moreover, these effects disappeared in a third sample of participants who received stroking to the palm. Yet, across all samples stroking velocity monotonically predicted EEG power in Rolandic rhythms and thus cortical activity as would be expected from the Aß processes known to be supported by somatosensory cortex (Case et al., 2016; Löken et al., 2009).

Together, these results imply that CT input, apart from traveling slowly to thalamus and insula, also takes a fast route to somatosensory cortex. Moreover, the temporal course of the observed effects points to interactions between slow unmyelinated and fast myelinated signaling in the spinal cord. Indeed, the present N400 effect emerges too early for a strictly CT driven response and too late for a strictly Aß driven response but converges with the timing of CT signals arriving in the dorsal horn. In detail, Aß projection velocities may be 70 m/s or faster (Susuki, 2010), whereas CT projection velocities are reported with a mean of 1 m/s and a maximum velocity of up to 2 m/s (Vallbo et al., 1993; Watkins et al., 2017). The mean distance between the forearm stimulation site and the cortex was ∼98 cm in a subset of 10 participants of the present study. The mean distance between the forearm stimulation site and the spinal cord was ∼57 cm. Thus, pure Aß input could reach cortex after ∼14 ms, while CT input could reach cortex after ∼490 ms (not taking into account small/unknown delays in cuneate and thalamic synaptic relays). Importantly, CT input would start to arrive in the spinal cord already after ∼285 ms, which is only slightly earlier than the 300 ms at which the present N400 effect developed.

Based on these figures, we speculate that CT projections to the dorsal horn interfaced with Aß pathways that then quickly forward CT information to the somatosensory cortex. This speculation should be corroborated in further human research by for example measuring individuals lacking C (Morrison et al., 2011) or Aß fibers (Olausson et al., 2002), but it does agree with extant non-human work. Research on the rodent LTMR recipient zone (Abraira et al., 2017) points to the existence of inhibitory and excitatory interneurons that merge input from Aß and C-LTMR fibers.

Additionally, there is evidence from cats that neurons in somatosensory cortex may show preferential responses to slow as opposed to fast stroking of hairy skin (McKenna et al., 1984). Our findings also converge with a recent study in human patients undergoing spinothalamic tract ablation (Marshall et al., 2019). Whereas the procedure impaired the perception of pain, temperature, and itch, it preserved the perception of pleasantness from the stroking of hairy skin. In light of the present results, this may be because the surgery preserved CT modulations of Aß projections via the dorsal-column medial lemniscus pathway. As we show here, these Aß projections can predict perceived pleasure and may temporarily suffice if CT input that accompanied them throughout a person’s lifetime is unavailable.

An interaction between Aß and C fiber pathways is central to the gating theory of pain perception (Melzack and Wall, 1965; Mendell, 2014). The present study suggests that a similar mechanism might be at play in the context of pleasurable touch. However, rather than Aß signals dampening the onward projection of C fiber input, it seems that C fiber input dynamically modulates Aß onward projections. As a result, somatosensory cortex can represent aspects of CT information before the original arrives. Such a fast-forwarding of CT information is likely to have consequences for ongoing brain and spinal processing. At the level of the brain, this could serve to prepare relevant somatosensory and socio-emotional networks for CT input. For example, it might modulate the sensitivity of insula and posterior superior temporal cortex for gentle caress (Morrison, 2016). Additionally, a cortical preview of CT signaling could have top-down influences on the activity of second order C fiber neurons in the spinal cord (Tinnermann et al., 2021, 2017). Depending on other ongoing processes, such somatosensory effects on brain and spinal networks could up or down-regulate CT representations and may constitute one possible mechanism by which contextual factors (e.g., relationship with a toucher) modulate tactile pleasure. The present study joins a couple of earlier attempts to investigate the temporal course of CT and Aß cortical responses. These earlier studies failed to report an N400 effect, but this may be because past designs probed only one (Ackerley et al., 2013; Hagberg et al., 2019) or two velocities (Haggarty et al., 2020) and were hence not set-up to observe velocity tuning. Indeed, the results we obtained readily replicated in a second sample, although not every tested individual produced a u-shaped N400 modulation—as one would expect given natural variation in a number of factors presumed to shape CT signaling (Lee Masson et al., 2019; Sailer and Ackerley, 2017). Yet, a quadratic pattern was discernible in the majority of participants and reached significance in 98 % of tests done for random sample subsets of only 15 participants (Supplementary Materials). Thus, the somatosensory N400 could proof useful in further research on affective touch. Specifically, this ERP effect could help elucidate stimulus (e.g., temperature), context (e.g., social partner), and person dependent aspects of CT processing. For example, one might examine whether childhood touch deprivation and autistic traits dampen or modify the somatosensory N400. Given that measuring CT activity in the periphery is very time consuming and only available to a handful of labs world-wide, a more accessible central marker should be of great utility for the field.

Before closing, we wish to highlight that the present study, by shedding light on the neuronal pathways involved in the perception of caress, builds on a number of methodological advances. Past studies relied on simple linear strokes that were typically applied with a rotatory motion device. Stimuli with different velocities also differed in duration (e.g., slower stimuli entailed longer durations). Moreover, variation in the curvature of arms and palms could not be accommodated such that stimuli tended to travel over short distances only. The present study tackled these issues using a novel tactile stimulation device with the same degrees of freedom as a hand touching skin. Thus, we could implement oval strokes, which are more natural (Lo et al., 2021) and deemed more pleasant than linear strokes (Shirato et al., 2018). We could also equate stimulus duration across conditions, which is critical when dealing with temporally sensitive measures such as the EEG. Moreover, we achieved this while controlling for skin site effects by counterbalancing the starting position of strokes across conditions. Last, our strokes readily traced a participant’s arm or palm curvature, thus allowing us more than twice the typical experimental stroking distance. We believe that future research will benefit from these and other modifications that make an experimental touch stimulus more natural.

To conclude, past research divided our tactile sense into a discriminative system served by Aß afferents and an affective system served by CT afferents. We offer here the first human evidence to challenge this divide. Our data help characterize a late frontal modulation that shows CT-like velocity tuning for input from both glabrous and hairy skin. Additionally, our data identify a novel somatosensory brain potential with CT-like velocity tuning that emerges well before C fiber signaling reaches the brain. Moreover, the temporal characteristics of this potential and its absence for glabrous stimulation point to spinal cord interactions between CT and Aß processes as the underlying mechanism. Thus, rather than depending on a single tactile system, the pleasure of caressing touch is caused by multiple systems negotiating the impact of bottom-up and top-down processes on an individual’s experience.

## Materials and Methods

### Resource Availability

Given the exploratory nature of Experiment 1, we first replicated key findings in another independent sample (Supplementary Materials) and, after successful replication, conducted a second experiment targeting glabrous, rather than hairy, skin. These efforts were not pre-registered as they immediately derived from the findings of Experiment 1. The study protocol was approved by the Clinical Research Ethics Committee of Hong Kong.

### Subject Details

We recruited 37 participants for Experiment 1, seven of whom were excluded from statistical analysis because noise in the EEG signal resulted in less than 30 epochs in one or more conditions when epoching was done using a 4 s window. However, six participants had sufficient data when epoching was done using a 2 s window, so their data were subsequently included in the replication of Experiment 1 as reported in the Supplementary Materials. The final sample of Experiment 1 comprised 15 men and 15 women with a mean age of 21 years (SD 2.98). We recruited 30 participants for Experiment 2 including 15 men and 15 women with a mean age of 20.1 years (SD 2.09). All participants reported being right-handed.

### Method Details

#### Stimuli and apparatus

The tactile stimuli were delivered using a custom-built tactile device capable of 3D motion (https://drive.google.com/file/d/1JclE_8DQvEef9NtKM6Xu0cYx72aF6J3F/view). The device was controlled via Matlab and entailed 8 motors that could move a touch stimulator in any direction with high spatial and temporal precision. In keeping with previous research, the touch stimulator in this study was a soft cosmetic brush. To enable temporal alignment between touch onset and the EEG, we weaved soft copper wires into the brush that connected with ESP32 Capacitive Touch Sensor pins. When the brush contacted skin, these pins sent a signal to the EEG data acquisition computer. The touch sensing pins also facilitated calibrating the touch device for a given participant. They enabled position read-outs for a planned stroking trajectory allowing us to adjust this trajectory to the surface curvature of the target skin area.

The touch stimuli in this study comprised gentle stroking along an oval trajectory. The set points for this trajectory were a ∼15 cm circumference, a minor radius of ∼1 cm and a major radius of ∼3.22 cm. Please note that small deviations from these set points were necessary due to variation in skin area curvature across participants. Strokes were delivered at five velocities including 0.5, 1, 3, 10 and 20 cm/s for a duration of 2.5 s. Because different velocities necessarily covered different distances across the skin, we adjusted the starting position of strokes such that motion along the oval was balanced across trials for a given velocity. Moreover, all velocities completed the full oval at least once. Note, however, that faster velocities stroked over the same area more frequently than slower velocities, as they had to travel larger distances (for further details, please see Supplementary Materials). In Experiment 1, the brush stroked the participant’s dorsal forearm, whereas in Experiment 2, the brush stroked her/his palm.

#### Procedure

After completing an informed consent procedure, the participant was seated and the experimenter prepared her/him for the EEG recording. The participant then placed her/his left forearm onto a comfortable arm rest under the touch stimulator. In Experiment 1, the arm was placed with a supine position, whereas in Experiment 2, it was placed with a prone position. As the prone position was slightly more effortful to maintain for the duration of the experiment, we attached a couple of soft elastic straps to gently assist the participant. A curtain precluded the participant from seeing the forearm and the touch device.

Next, the participant received instructions over a computer monitor placed in front of her/him. The participant was asked to insert noise-canceling ear-phones into the ears which presented a soft white noise meant to block out any remaining noise from the movement of the touch device. Then, the experimenter operated the device and re-adjusted the white noise volume until the participant no longer heard the touch device. After ensuring the participants’ comfort, the experimenter started the experiment.

The experiment comprised 300 trials in which the five stimulus velocities were presented with equal probability in pseudo-random order such that the same velocity would not be presented consecutively. A trial began with a fixation cross lasting for 0.4 to 0.55 s coinciding with the downward motion of the touch stimulator. After the stimulator contacted the skin, it began moving along the oval trajectory for 2.5 s. During this time and the following one second, the fixation cross remained on the computer screen and was then replaced by a pleasantness rating scale. The participants now used their right arm to operate a mouse and to move a cursor to a position on a continuous scale that reflected the pleasantness associated with the touch. The scale endpoints were marked with very unpleasant on the left and with very pleasant on the right and scores coded within a range of -100 to 100. Following the participant’s response, there was a short inter-trial interval during which the screen remained blank. The interval lasted for 1, 1.5 or 2 seconds, drawn from a uniform distribution. The experiment was divided into four blocks of 75 trials. Participants had a short break after every 30 trials and a five-minute break between each block. Trials lasted about 6.5 seconds and an experimental session lasted about 50 minutes.

#### Electrophysiological recording

The EEG was recorded using 64 Ag/AgCl electrodes, which were located according to the extended 10–20 system of the American Clinical Neurophysiology Society (Acharya et al., 2016). CPz was used as the online reference. Electrode impedance was below 20 kΩ. The data were recorded at 500 Hz with a BrainAmp EEG system. Only an anti-aliasing filter was applied during data acquisition (i.e., sinc filter with a half-power cut-off at half the sampling rate).

### Quantification and statistical analysis

EEG data were pre-processed with EEGLAB v14.1.1 (Delorme and Makeig, 2004) implemented in MATLAB. The data were down-sampled to 250 Hz, low-pass filtered at 30 Hz (7.5 Hz transition bandwidth, -6 dB cut-off) and high-pass filtered at 0.1 Hz (0.1 Hz transition bandwidth, -6 dB cut-off). Then the data were re-referenced to the channel average and epoched with a window from -1 to 3 s around each stimulus onset (Experiment 1) or from -1 to 1 s around each stimulus onset (Experiment 1 & 2). Afterwards, the data were subjected to manual inspection where channels and epochs with non-typical artifacts caused, for example, by muscle movements or drifting were rejected. The cleaned data was then high-pass filtered at 1 Hz and subsequently entered in an adaptive mixture independent component analysis (AMICA) (Palmer et al., 2011). The resulting independent component structure was applied to the original data with the 0.1 - 30 Hz filter setting. Components reflecting typical artifacts (i.e., horizontal and vertical eye movements and eye blinks) were removed and the data was back-projected from component space into EEG channel space. The data were subjected to another round of visual inspection during which residual artifacts were removed. A current source density transformation was applied using the CSD Toolbox (Kayser and Tenke, 2015). This served to enhance spatial separation of temporally overlapping signal components and to facilitate the detection of independent cortical sources (Kamarajan et al., 2015).

For the ERP analysis, we then conducted a baseline correction using mean voltages within a window between -200 and 0 ms from stimulus onset. Subsequently, trial data were subjected to a subject and condition-wise averaging routine. For the time-frequency analysis, we subjected epochs ranging from -1 to 1 s to a continuous wavelet transformation with cycles ranging from 3 to 7 for frequencies from 5 to 28 Hz in steps of 1 Hz. This returned 153 time points ranging from -663 and 667 ms around stimulus onset. The wavelet transforms were then baseline corrected using a window from -500 to -100 ms and their power was obtained and averaged for each participant, condition, time point, and frequency.

Ultimate trial numbers per condition averaged across participants ranged from 50 to 53 for the long epochs in Experiment 1 (participant-wise min = 31 and max = 68). They ranged from 55 to 57 for the short epochs in Experiment 1 (participant-wise min = 37 and max = 74) and from 54 to 56 for the short epochs in Experiment 2 (participant-wise min = 37 and max = 69).

