## Supplementary Materials for "Discriminative and affective touch converge: Somatosensory cortex represents Aß input in a CT-like manner"

### Experiment 1

#### *Additional Method Details*

Table 1. Velocity condition information

| Velocity (cm/s) | Stroking Time (s) | Stroking Distance (cm) | Distance larger than the total stroking circumference? | Required Trials |
|---|---|---|---|---|
| 0.5 | 2.5 | 1.25 | No | 12 |
| 1 | 2.5 | 2.5 | No | 6 |
| 3 | 2.5 | 7.5 | No | 2 |
| 10 | 2.5 | 25 | Yes | 6 |
| 20 | 2.5 | 50 | Yes | 6 |

Note. Five stroking velocities of 0.5, 1, 3, 10, 20 cm/s were tested. The stroking time was kept constant such that the stroking distance varied across conditions. “Required Trials” refers to the number of trials needed to complete one stroking circumference and determined within-subject counterbalancing. For the velocities of 0.5 cm/s, 1 cm/s, and 3 cm/s, the stroking distance on a single trial will be less than the full circumference of the stroking trajectory (15 cm). The required trials will be the number of times needed to achieve 15 cm. For example, trials of 0.5cm/s only cover 1.25 cm in stroking distance thus necessitating 12 trials to complete the 15 cm long elliptical trajectory. For the velocities of 10 cm/s and 20 cm/s, the stroking distance on a single trial will be greater than the defined circumference but cannot be perfectly divided by 15 cm. Without altering the starting positions, some locations on the arm would be stroked more frequently than the others. To control this, we adjusted the onset position along the defined circumference.

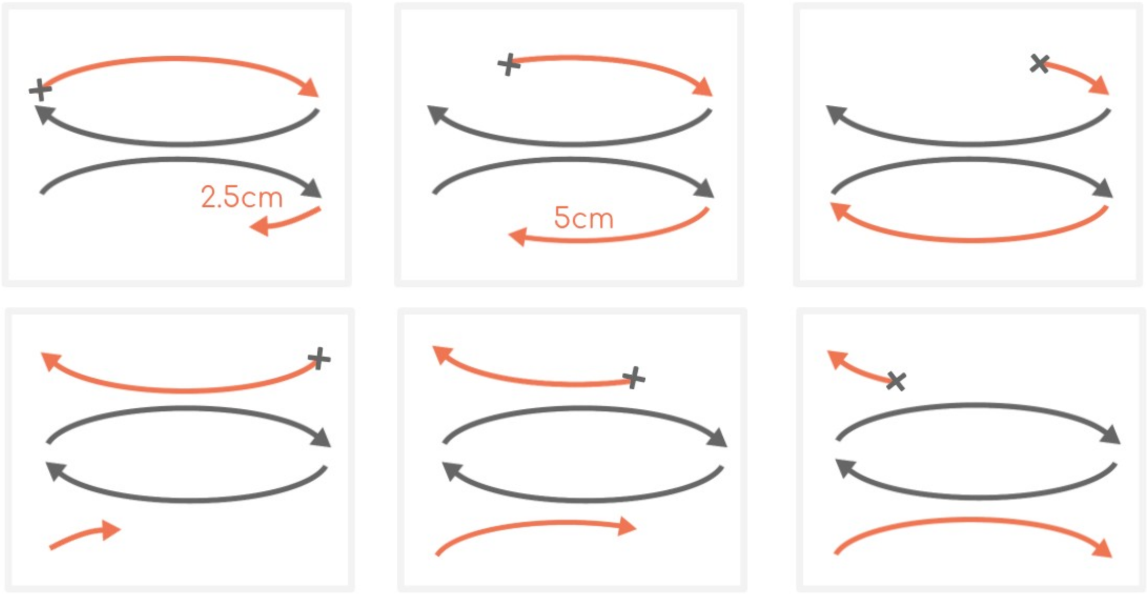

Figure 1. Exemplary counterbalancing of touch onset positions for stroking velocities of 10 cm/s. Each graph shows one of six touch onset positions counterbalanced within participants. As the stroking distance was 25 cm, there were 10 cm excess after one complete ellipse trajectory (15 cm). To distribute this excess distance evenly across the whole ellipse, we changed the starting position in steps of 2.5 cm. Black arrows indicate a complete elliptical trajectory; red arrows indicate the excess distance. The cross indicates the starting position. The concatenation of all the red arrows in the six motion patterns forms three complete elliptical trajectories.

***Subject-wise N400 plots***

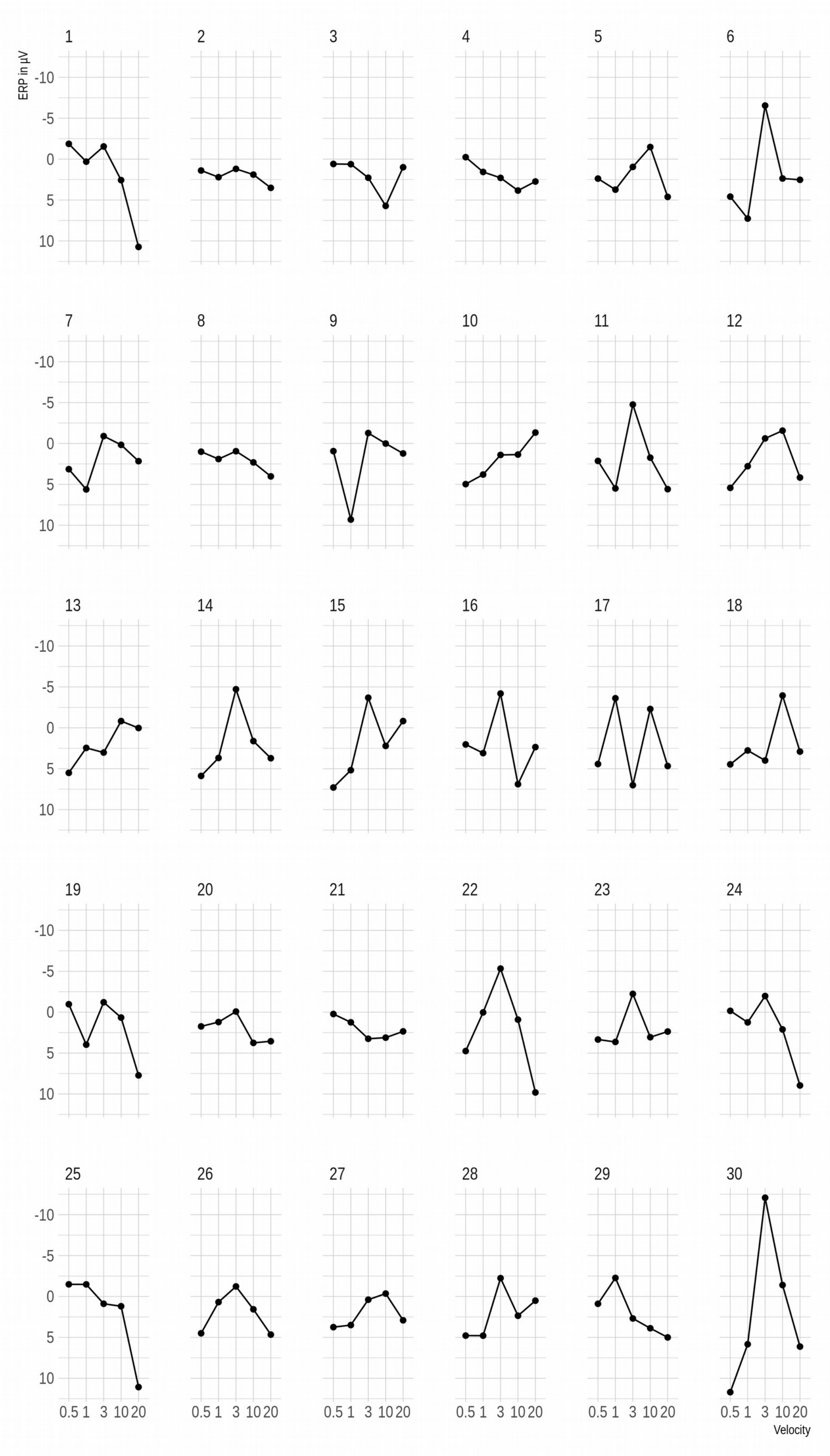
ERP in µV
Velocity
-10
-5
0
5
10
0.5 1 3 10 20

Figure 2. Mean voltages recorded over the right medial somatosensory cortex (Cz, C2, CP2) contra-lateral to the stimulation site. Each dot represents the ERP averaged between 300 and 500 ms following stimulus onset. Although not all participants clearly show an inverted u-shaped relationship between the ERP and velocity, most participants do. As such, this component might be useful for studying inter-individual differences in CT signaling and sensititive to "affective touch".

### Experiment 1 - Replication

#### *Methods*

*Participants*. We recruited 31 participants for this experiment. Due to technical issues, one data set had to be excluded. The final data came from 15 men and 15 women with a mean age of 21.2 years (SD 2.51). All participants were right-handed.

*Materials and Procedure.* Materials and procedure were identical to Experiment 1 reported in the main body of the manuscript.

#### *Results*

*ERPs.* The results are illustrated in Figure 2. As for Experiment 1, we computed Fz mean voltages from 700 ms to 1000 ms, the end of the stimulus epoch and subjected those means to a mixed effect model with the second order polynomial of the natural velocity logarithm as the fixed effect and the participants' intercept as the random effect. This revealed a significant velocity effect ($F[2,118]=5.78$, p=.004). Additionally, a likelihood ratio test with a model without the polynomial indicated that the polynomial offered a better fit (X[1]=20.93, p<.0001).

We again observed a modulation between 300 to 500 ms following touch onset over somatosensory leads. Thus, we computed mean voltages within this time window and across the same right centro-medial electrodes (Cz, C2, CP2) used in Experiment 1. A polynomial mixed effect model was again significant (F[2,118]=20.53, p<.001) and was superior when compared with a simple linear model (X[1]=43.44, p<.0001).

*Rolandic rhythms over contra-lateral somatosensory cortex.* Like the ERP, mean power in the alpha and beta band at right centro-medial electrodes was subjected to a polynomial mixed effects model with significant results (F[2,118]=12.15, p<.001). Note, however, that the quadratic term indexed a monotonical decrease in power with increasing velocity. Again, the polynomial fit was superior over a linear fit (X[1]=4.37, p=.036).

*Pleasantness Ratings.* Rating data were analyzed as a function of the natural logarithm of velocity using a mixed modeling approach. Subject and trial wise rating scores were subjected to a model with the second order polynomial of the velocity variable as the fixed effect and the participants' intercepts and slopes as the random effects. F-statistics were obtained using the Satterthwaite approximation for degrees of freedom. This revealed a significant velocity effect (F[2,29]=6.44, p=.005). Additionally, a likelihood ratio test comparing this original model against one with velocity modeled linearly revealed that the former was a significantly better fit (X[4]=1413.5, p<.0001).

Next, we examined which brain measures predict pleasantness ratings. Pleasantness served as the dependent variable, a given brain measure and velocity served as fixed effects, and the participants' intercepts served as the random effect. Velocity was included simply to reduce Type 1 error and was of no interest here. For the Fz ERP, the result was non-significant (p=.692). However, we observed again a significant relationship between the somatosensory N400 (F[1,147]=6.27, p=.013) indicating that more negative ERPs were associated with greater pleasantness. The power of mu rhythms over the somatosensory region showed a marginally positive association with pleasantness (F[1,293]=3.79, p=.052).

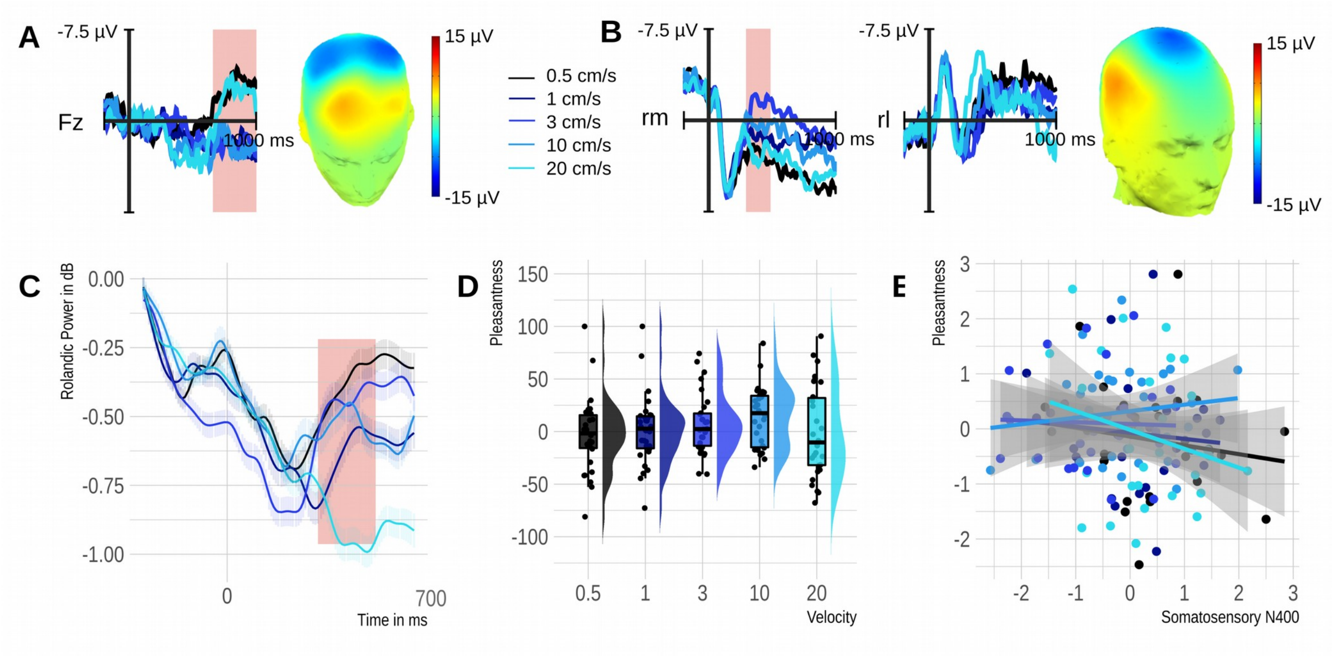

Figure 3. Results of the Experiment 1 Replication. A) Velocity effect observed at Fz. B) Somatosensory N400 velocity effect over the right medial region explored in Experiment 1 and its polarity inversion over the right lateral region. Reddish rectangle highlights windows for statistical analysis ranging from 0.7 to 1 seconds for the Fz effect and from 0.3 to 0.5 seconds for the somatosensory N400. The topography of each velocity effect (3 cm/s minus 20 cm/s) is printed on a 3D head. Fz – frontal midline electrode, rm – right medial region of interest, rl – right lateral. C) Time course of power changes evoked by tactile stimuli in alpha and beta bands. D) Rain plots illustrate a

change in subjective pleasantness as a function of stimulus velocity. E) Scatter plots with regression lines showing the linear relationship between the normalized subjective pleasantness and the normalized N400 amplitude at the subject level. Overall, greater pleasantness was associated with a larger (more negative) N400 amplitude.

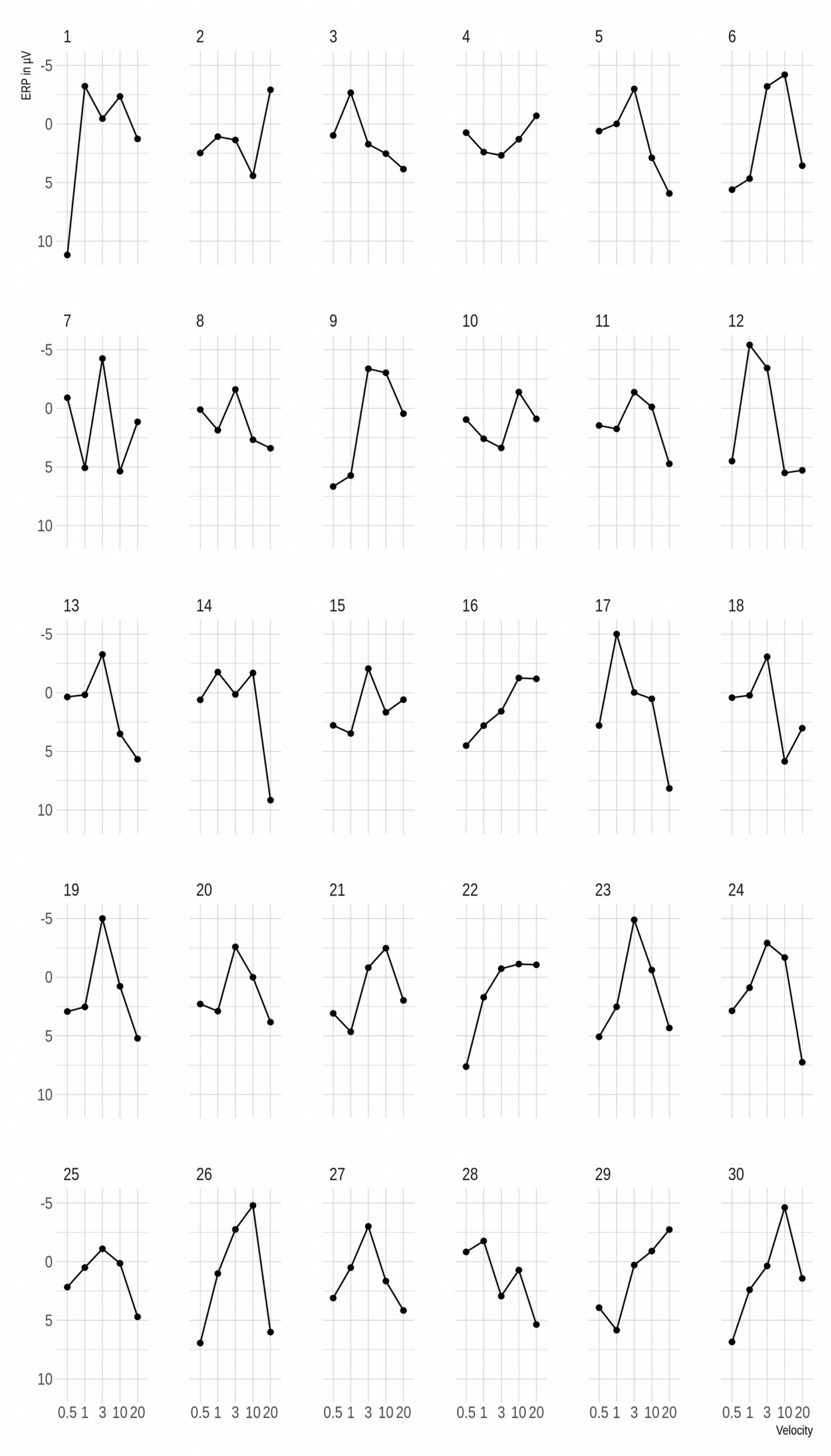
ERP in µV
Velocity
-5
0
5
10
0.5 1 3 10 20

Figure 4. Mean voltages recorded over the right medial somatosensory cortex (Cz, C2, CP2) contra-lateral to the stimulation site. Each dot represents the ERP averaged between 300 and 500 ms following stimulus onset.

### Permutation based testing of the N400 in a smaller sample

Individual data plots of the somatosensory N400 implied that the effect is relatively robust and shows for the majority of participants. To explore how many participants may be needed to obtain a statistically significant quadratic N400 effect, we pooled the data from Experiment 1 and its replication. We then randomly selected (without replacement) 10 or 15 samples for analysis. Again the data from these 10 samples entered a mixed effect model with the second order polynomial of the natural logarithm of Velocity as the fixed effect and the participants' intercepts as the random effect. This procedure was repeated 5000 times and the number of significant tests ($p<.05$) divided by the number of all tests. The results revealed a probability of 0.87 for samples of 10 participants and a probability of 0.98 for samples of 15 participants.
